# Catch me if you can: current status and topical issues on the use of eDNA-based targeted detection of rare and endangered animal species

**DOI:** 10.1101/2023.06.13.544783

**Authors:** Sofia Duarte, Luara Simões, Filipe O. Costa

**Affiliations:** Centre of Molecular and Environmental Biology (CBMA) and ARNET-Aquatic Research Network, Department of Biology, University of Minho, Campus de Gualtar, 4710-057 Braga, Portugal; Institute of Science and Innovation for Bio-Sustainability (IB-S), University of Minho, Campus de Gualtar, 4710-057, Braga, Portugal

**Keywords:** eDNA-based tools, endangered and invasive species, species-specific assays, environmental factors effects

## Abstract

Animal detection through DNA present in environmental samples (eDNA) is a valuable tool for detecting rare species, that are difficult to observe and monitor. eDNA-based tools are underpinned by molecular evolutionary principles, which are key to devising tools to efficiently single out a targeted species from an environmental sample, using carefully chosen marker regions and customized primers. Here, we present a comprehensive review of the use of eDNA-based methods for the detection of targeted animal species, such as rare, endangered, or invasive species, through the analysis of 460 publications (2008-2022). Aquatic ecosystems have been the most surveyed, in particular, freshwaters (75%), and to a less extent marine (14%) and terrestrial systems (10%). Vertebrates, in particular, fish (38%), and endangered species, have been the most focused in these studies, and Cytb and COI are the most employed markers. Among invertebrates, assays have been mainly designed for Mollusca and Crustacea species (22%), in particular, to target invasive species, and COI has been the most employed marker. Targeted molecular approaches, in particular qPCR, have been the most adopted (73%), while eDNA metabarcoding has been rarely used to target single or few species (approx. 5%). However, less attention has been given in these studies to the effects of environmental factors on the amount of shed DNA, the differential amount of shed DNA among species, or the sensitivity of the markers developed, which may impact the design of the assays, particularly to warrant the required detection level and avoid false negatives and positives. The accuracy of the assays will also depend on the availability of genetic data from closely related species to assess both marker and primers’ specificity. In addition, eDNA-based assays developed for a particular species may have to be refined taking into account site-specific populations, as well as any intraspecific variation.

**Graphical Abstract:** 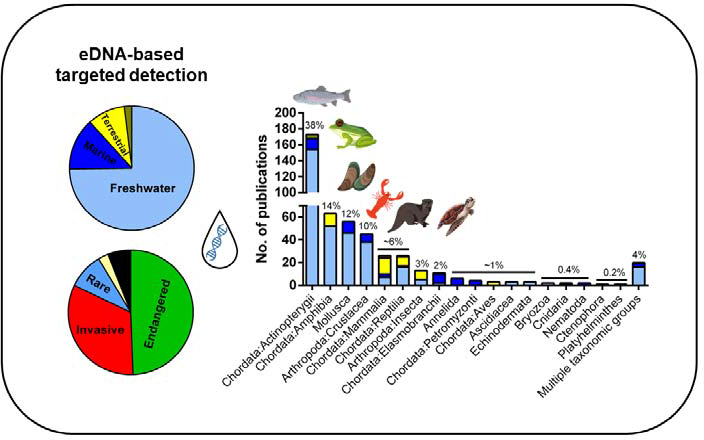

## 1. Introduction

Biodiversity plays a key role in maintaining the integrity of ecosystems, by providing important services such as buffering extreme climate events, regulating hydrological cycles and temperature in urban areas, protecting soils, economic diversification, and reducing food insecurity (Naeem et al., 2016). Biodiversity monitoring is, thus, essential for assessing ecosystem health, in particular, to signal endemic endangered species or to early detect invasive species, which are all crucial to provide guidelines for more effective management of natural resources (Navarro et al., 2017).

Species monitoring has been relying customarily on species visualization or capture and identification of specimens through diagnostic morphological characters (Qu and Stewart, 2019). For instance, aquatic vertebrates’ surveillance programs traditionally emplsoy nets or electrofishing gear (Goldberg et al., 2011; Jerde et al., 2011). However, this is a process that can be laborious, time-consuming, and deficient in taxonomic discrimination capacity, in particular for organisms low on distinctive morphological features, such as the case of some invertebrate fauna. In addition, when the targets are rare species, i.e., species with small populations sizes, or elusive species, the detection probabilities are typically low in any ecosystem and a greater sampling effort is needed to maximize the chances of species detection, which is not always feasible (Delgado, 2022; Goldberg et al., 2011; Jerde et al., 2011; Ma et al., 2022; Sgarbi et al., 2020). Although the term “rare species” is commonly associated with indigenous endangered species, non-indigenous species can be considered rare species as well, namely early in the invasion process when their population sizes are still small (Ficetola et al., 2008; Goldberg et al., 2011; Jerde et al., 2011).

The adoption of more sensitive and non-invasive methods, with higher detection capacity, can be particularly advantageous in the case of rare species detection. Methods that rely on the use of environmental DNA (eDNA) have been placed at the forefront for their great potential in biodiversity monitoring (Ficetola et al., 2008; Jerde et al., 2011; Leese et al., 2016, 2018). In addition, contrarily to customary invasive approaches, eDNA sampling minimizes or avoids any disturbance to the target and co-occurring species and sampled habitat. Environmental DNA is obtained directly from environmental samples (i.e., water, soil, sediments, air) and can exist mainly in two forms: either contained in cells of small organisms such as microbes, single-celled algae, meiofauna, and zooplankton or in free-form that is released by larger organisms into the environment through faeces, urine, mucous, gametes, skin cells, among other particles (Pawlowski et al., 2020; Thomsen et al., 2012). Once eDNA is shed into the environment, its persistence may vary from hours to weeks in temperate waters, to several months or years in soil, caves, permafrost, or sediments (Baillie et al., 2019; Barnes et al., 2014). However, eDNA is still presumed to be the predominant source of organismal DNA and indicative of the organism’s recent presence, but it can be also highly dependent on the system under analysis (Thomsen and Willerslev, 2015).

Species detection through eDNA has been mainly achieved using two following approaches: i) targeted species detection or active surveillance, where specific primers are used for detection of single or few species using a PCR platform (Ficetola et al., 2008; Goldberg et al., 2011; Jerde et al., 2011; Wood et al., 2019b) and ii) community-level detection, or passive surveillance, where a complete inventory of the species, within a given ecosystem or habitat, is accomplished using either broad-spectrum or taxonomic groupLspecific primers, in combination with high throughput sequencing (HTS), .i.e., eDNA metabarcoding (Taberlet et al., 2012a, 2012b). In the latter case, abundant, rare, endangered, and invasive species or the diversity of a specific taxon (e.g., fish) will be concurrently assessed. The high sensitivity of eDNA-based detection and the greater probability of tracking rare species in their habitat, typically results in higher species richness estimates, associated with lower sampling costs and survey times, compared with classical surveys (Belle et al., 2019; Coble et al., 2019; Pawlowski et al., 2018; Taberlet et al., 2012a, 2012b; Xia et al., 2021). In addition, species that are present even at low abundances, such as the case of rare species, can be efficiently detected (Thomsen and Willerslev, 2015). While the application of eDNA metabarcoding is increasing significantly, whereas studies applying single species detection are declining (Schenekar, 2023), targeted species detection is still the best strategic approach for detecting one or a few species at a specific location and time, increasing the likelihood of detection (Morisette et al., 2020). In addition, targeted detection is particularly advantageous for “finding the needle in the haystack” and when the target is of high risk if it goes undetected (Harper et al., 2018b; Morisette et al., 2020). For instance, to detect an invasive species early in the invasion process, knowing the characteristics of the target species and being in the right place, at the right time, and using the most appropriate tools increases the chances of successfully addressing a bioinvasion, in an effective and cost-efficient manner (Morisette et al., 2020). In addition, when using community-level assessments such as eDNA metabarcoding, the co-occurrence of abundant species can reduce the probabilities of rare species detection (Gargan et al., 2022; Harper et al., 2018b; Rojahn et al., 2021). Since the pioneer studies of Ficetola (2008), Goldberg (2011), Jerde and co-authors (2011), among others, consisting on the targeted detection of invasive and rare animal species, that the employment of eDNA-based tools for monitoring species of conservation interest has been rising dramatically (Belle et al., 2019; Bohmann et al., 2014; Rees et al., 2014; Thomsen et al., 2012). Thus, given the considerable amount of information already existing, an appraisal of the use of eDNA-based targeted detection is timely and much needed. To that end, we conducted a comprehensive review to analyse what geographic regions and ecosystems have been mostly surveyed, the taxonomic groups that have been targeted and species status (i.e., endangered or invasive), the platforms employed, as well as the DNA markers and length, and assays that have been already implemented by environmental managers to support conservation-related decisions. In addition, we also assessed potential gaps and biases, as well as the greatest challenges that are still to be addressed, and recommend future development in the context of biological conservation.

## 2. Methods

We performed a literature search by querying the Web of Science for articles in which eDNA-based tools were used for detecting rare species, on July 1^st^, 2022. The search was limited to titles, abstracts, and keywords (search by topic), that contained the terms “environmental DNA” OR “eDNA” and terms categorizing the target groups, namely “rare” OR “elusive” OR “endangered” OR “threatened” OR “imperiled” OR “vulnerable” OR “invasive”. The search retrieved 1,049 articles, published between 2006 and 2022 (until 30^th^ June) (**Table S1**). After individual inspection, we retained 460 articles for conducting our analysis, published between 2008 and 2022 (**Table S2**). Papers that were not primary research articles (e.g., reviews) or surveyed the biodiversity of whole communities using eDNA metabarcoding were excluded from the analysis. Furthermore, studies were included in the analysis only if they specified that the aim was to target one or a few rare/elusive wild species. Since our survey is focused on animals, studies aiming at other taxonomic groups such as plants, fungi, bacteria, and protists, among others, were also excluded.

From each selected publication we retrieved the following information: i) the geographic area/country, ii) the environment (e.g., terrestrial, freshwater, and marine), iii) the type of environmental sample (e.g., water, sediment, soil, among others), iv) if the study was conducted in the field or a controlled environment (e.g., aquarium, mesocosms, lab), v) the targeted species, the respective taxonomic classification and species category (i.e., endangered, invasive), vi) the targeted molecular markers and segments length (bp) and vii) the platforms employed (e.g. cPCR, qPCR) (**Table S2**). We did not analyse in detail all protocols used through the analytical chain of eDNA-based targeted detection (i.e., sampling, eDNA capture, eDNA extraction protocols), since these have been already the target of previous reviews (Doi et al., 2021; Kumar et al., 2020; Lear et al., 2018; Rees et al., 2014; Shu et al., 2020; Tsuji et al., 2019; Wang et al., 2021). Linear regressions were used to assess the significance of the increase in the number of papers per year, between the periods of 2008 and 2015 and the periods of 2016 and 2022, using GraphPad Prism v6 (GraphPad Software, Inc.).

## 3. Results and Discussion

Most of the original publications retrieved from the initial list belonged to the categories of Ecology, Biodiversity Conservation, Environmental Sciences, and Marine and Freshwater Biology (**Fig. S1**). A detailed analysis of the 460 articles retained, indicated that papers addressing the use of eDNA-based tools for the targeted detection of rare animal species have been published in 124 scientific journals (**Table S2**), but only 25 journals have been selected in more than 1% of the publications (at least 5 publications) (**Fig. S2**). The most frequently selected journal was PLoS ONE (54 publications), followed by Conservation Genetics Resources (27) and Biological Invasions (22) (**Fig. S2, Table S2**).

Since the pioneer study of Ficetola and co-authors (Ficetola et al., 2008), which developed a species-specific assay for detecting the invasive bullfrog using water eDNA collected in French wetlands, the number of publications addressing eDNA-targeted detection in biological conservation studies has grown rapidly, with a particularly steep trend from 2016 onwards (Fig. 1). Indeed, the number of publications increased at a rate of 7.2 papers/year between 2008 and 2015 (P=0.04, R^2^=0.71), while a rate of 61.7 papers/year was found between 2016 and 2022 (P<0.0001, R^2^=1.0) (Fig. 1), which is congruent with the increasing adoption of eDNA-based tools in the last few years (e.g. Minamoto, 2022; Nordstrom et al., 2022; Schenekar, 2023).

**Figure 1.**
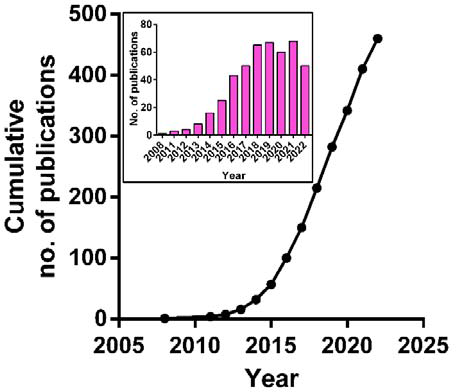
Total number of publications per year (insert) and the cumulative number of publications using eDNA-based targeted detection of rare animal species (n=460).

### 3.1. eDNA-based targeted detection of rare animal species has been applied mainly in aquatic ecosystems of North America and Europe

Most of the studies analysed in the current review have been conducted in a single geographic region targeting species with restrictive distributions (e.g. Fukumoto et al., 2015; Neice and McRae, 2021; Padgett-Stewart et al., 2016; Westhoff et al., 2022; Wilcox et al., 2014) (Fig. 2A, **Table S2**). A few have been conducted in multiple countries, but within a single continent (Agersnap et al., 2017; De Ventura et al., 2017; Rusch et al., 2020; Schneider et al., 2016; Thomsen et al., 2012), while very few have been intercontinental (Meekan et al., 2017; Ribani et al., 2020; Takeuchi et al., 2019). Most of the studies have been performed in North America (47%) and Europe (23%), demonstrating a bias towards the northern hemisphere, in what respects the adoption of these tools (Fig. 2B, **Table S2**). The geographic bias towards the Northern hemisphere has been previously documented before (Belle et al., 2019; Coble et al., 2019; Duarte et al., 2021a, 2021b; Nordstrom et al., 2022; Schenekar, 2023).

It has been pointed out that the lack of studies in the southern hemisphere is mostly due to socioeconomic constraints and a lack of supporting infrastructures required for eDNA-based monitoring implementation in global South countries, in particular in African countries (Belle et al., 2019; Schenekar, 2023). In addition, the higher number of existing legal frameworks in Northern regions such as the Water Framework Directive (WFD) (Council Directive 2000/60/EC) and Habitats Directive in Europe (Council Directive 92/43/EEC) or the Endangered Species Act and National Invasive Species Act in North America, requiring regular biomonitoring programs, may also help to explain the higher adoption of eDNA-based tools in these geographic regions.

Aquatic ecosystems have been the most surveyed, in particular, freshwaters (75%), and to a less extent marine (14%) and terrestrial systems (10%) (Fig. 2C), a pattern that has been found in the different geographic regions surveyed, with some few exceptions (e.g., in North and South America and Africa, terrestrial ecosystems have been more surveyed than marine ecosystems), while multiple typologies of ecosystems have been surveyed only in approximately 2% of the studies, e.g., freshwaters and marine (Kasai et al., 2020; Lehman et al., 2022); marine and terrestrial (Farrell et al., 2022; Steinmetz et al., 2021) (Fig. 2). This is not surprising since freshwater ecosystems are considered the most imperilled habitats in the world (Reid et al., 2019), and at least in temperate regions they have been subject of intensive monitoring due to legal requirements, as already above-mentioned. In these ecosystems, most studies have been conducted in rivers and small streams (Castañeda et al., 2020; Ma et al., 2016; Mizumoto et al., 2020; Piggott, 2017; Riaz et al., 2020; Rodgers et al., 2020), lentic ponds (Adams et al., 2019; Geerts et al., 2018; Harper et al., 2018b) and lakes (Johansson et al., 2020; Kamoroff and Goldberg, 2018), to a small extent in reservoirs and dams (Nakao et al., 2023; Sepulveda et al., 2022, 2019), channels (Beauclerc et al., 2019; Díaz-Ferguson, 2014), caves and springs (DiStefano et al., 2020; Vörös et al., 2017) and aquacultures (Deutschmann et al., 2019; Ladell et al., 2019) (**Table S2**).

Although marine ecosystems and diversity are also under threat, in comparison to freshwaters these have been much less targeted, in part due to their vastness and inaccessibility, and high complexity, which made the targeted detection of rare species using eDNA highly challenging to implement (Suarez-Bregua et al., 2022). In addition, dedicated studies are still needed to understand how environmental factors (e.g., temperature, currents, tides, depth, stratification, and salinity) affect eDNA distribution and persistence dynamics in the marine environment, to optimize/support both sampling and results interpretation (Collins et al., 2018; Suarez-Bregua et al., 2022). In marine ecosystems, most studies have been conducted in coastal areas or estuaries (Crane et al., 2021; Ellis et al., 2022; Miralles et al., 2019, 2016; Yip et al., 2021), coastal lagoons (Ardura et al., 2017; Muñoz-Colmenero et al., 2018), aquacultures (Brand et al., 2022; Matejusova et al., 2021) and harbours and recreational marinas (Kim et al., 2018; Matejusova et al., 2021; Wood et al., 2017), whereas fewer have been conducted in the open sea (Catanese et al., 2022; Gargan et al., 2022; Wada et al., 2020) (**Table S2**).

Despite eDNA-based monitoring has been highly used in terrestrial ecosystems for assessing soil microbial communities through metabarcoding, studies are scarce on what concerns eDNA-based targeted detection of rare animal species, as indicated by our review. The surveyed habitats were variable and included farms (Macgregor et al., 2021; Maslo et al., 2017), temporary wetlands (Feist et al., 2022; Schumer et al., 2019; Tarof et al., 2021), soil ecosystems (Kucherenko et al., 2018; Yasashimoto et al., 2021), bromeliads tanks or tree holes (Barata, 2021; Mullin et al., 2022; Torresdal et al., 2017), among others e.g., honey samples (Utzeri et al., 2021), roadside drains (Smart et al., 2015) (**Table S2**).

**Figure 2.**
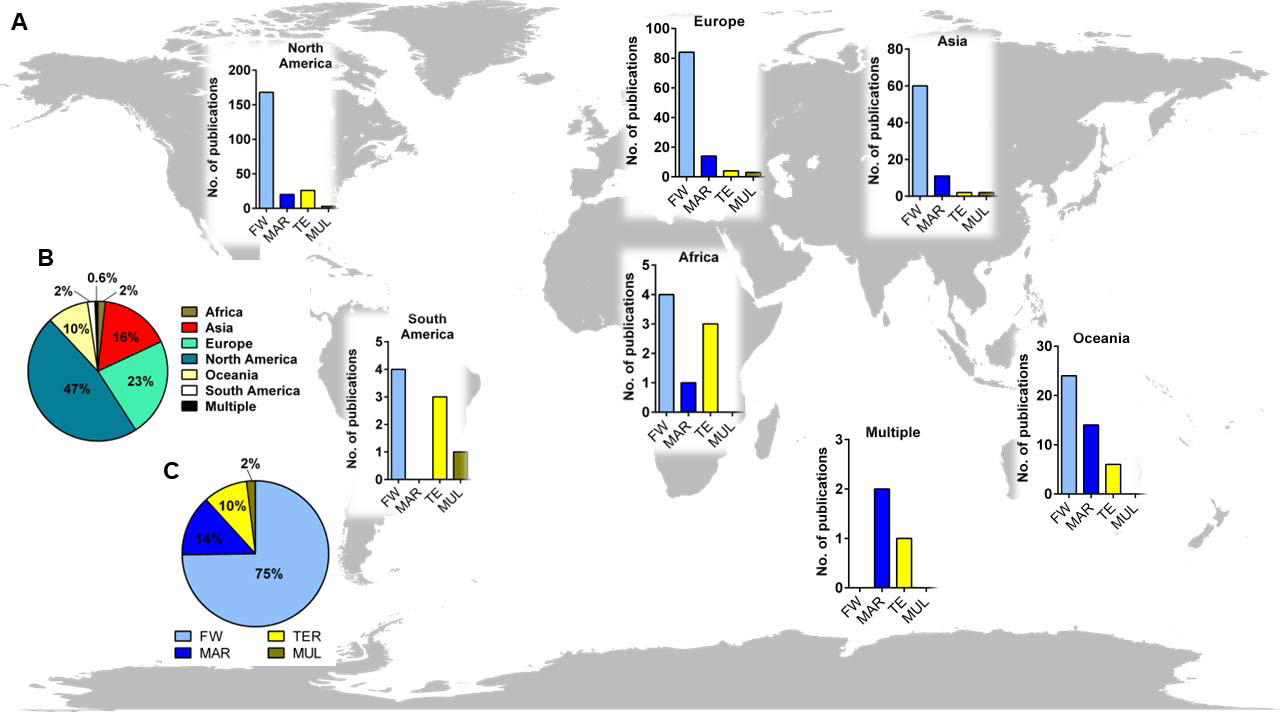
Geographic distribution of the 460 studies, separated by the environment surveyed (A) and geographic regions (B) and environments surveyed in the 460 publications (C), in %. FW: freshwater ecosystems, MAR: marine ecosystems, TER: terrestrial ecosystems, MUL: multiple ecosystems.

Water has been by far the most sampled in all types of surveyed ecosystems (>80% of the samples), as a result of most of the assays being designed for aquatic or semi-aquatic species (Fig. 3, **Table S2**). In water, eDNA can be distributed both dissolved or attached to suspended particles, which can eventually settle down into sediment layers. However, sediments have been less used as a source of eDNA (approx. 3% of the studies, Fig. 3). In terrestrial ecosystems it is difficult to find equivalent substrate types that effectively can capture rare animal species on land. In these ecosystems, soil (1.6%) (Matthias et al., 2021; Neice and McRae, 2021; Yasashimoto et al., 2021), faeces and urine (2.6%) (Steinmetz et al., 2021; Walker et al., 2017) or surfaces (2.4%), such as from plants or traps (Butterwort et al., 2022; Feist et al., 2022; Valentin et al., 2020), have been sampled as eDNA sources (Fig. 3). Other less used substrates include gut contents (Keskin, 2016), blood from invertebrates feeding on vertebrate species (e.g., leech blood parasiting turtles) (Farrell et al., 2022), air (Serrao et al., 2021) or tracks samples (Franklin et al., 2019).

**Figure 3.**
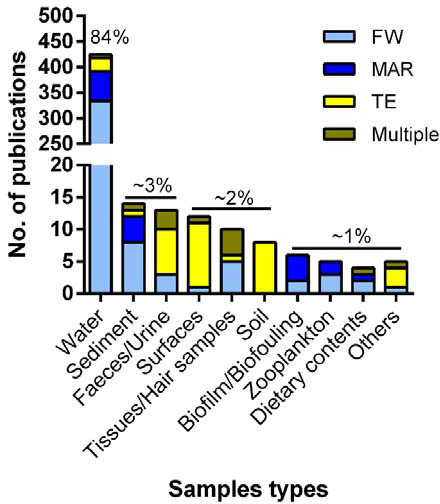
Types of samples used in the 460 publications using targeted assays for detecting rare animal species separated by the environment surveyed and % of each sample type used in the total number of publications (numbers above bars). FW: freshwater ecosystems, MAR: marine ecosystems, TER: terrestrial ecosystems, MUL: multiple ecosystems.

### 3.2. Fish and endangered species have been the main focus of eDNA-based targeted detection

Among the 460 studies, the taxonomic group for which the majority of targeted detection assays have been designed or employed is Chordata (approx. 67%) (Fig. 4). The targeted assays within Chordata have predominantly been adopted for the classes of Actinopterygii (38%) and Amphibia (14%) (Fig. 4). This is not surprising since a taxonomic bias towards fish has been previously observed in freshwater eDNA research (Belle et al., 2019), due to the high socioeconomic value of most species, which include both globally invasive and endangered species. In comparison, although the importance of several groups in biomonitoring (e.g., Arthropoda, Mollusca, and Annelida) or as invasive species or pests, invertebrates have been much less targeted using eDNA-based specific assays. Among invertebrate fauna, assays have been mainly designed for Mollusca (approx. 12%) and Arthropoda: Crustacea (approx. 10%). In freshwaters, a previous study on 272 peerLreviewed articles, published between 2005 and 2018, revealed that the targets of eDNA research about aquatic conservation were dominated by fish, followed by amphibians and molluscs, while freshwater arthropods were underLrepresented in their estimated species richness (Belle et al., 2019), which corroborates well with the results found in the current review. These findings are also in concordance with the fact that the investment per species is much higher for vertebrates in comparison to invertebrates, for example, in LIFE projects (EU’s funding instrument for the environment and climate action) (Mammola et al., 2020).

**Figure 4.**
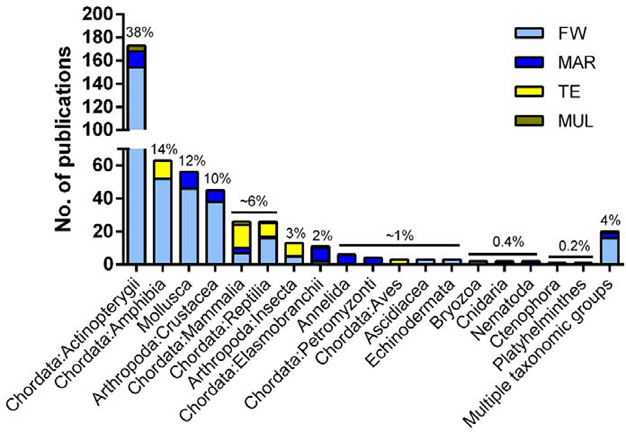
Taxonomic groups for which the targeted detection assays have been designed or employed in the 460 studies for detecting rare animal species, separated by the environment surveyed and % of each sample type used in the total number of publications (numbers above bars). FW: freshwater ecosystems, MAR: marine ecosystems, TER: terrestrial ecosystems, MUL: multiple ecosystems.

Most studies have designed assays for the detection of species within one single phylum (>95%), e.g., Chordata: Amphibia (Fukumoto et al., 2015; McKee et al., 2015); Chordata: Actinopterygii (Clusa and García-Vázquez, 2018; Jerde et al., 2013; Roy et al., 2018); Mollusca (Cho et al., 2016); Arthropoda: Crustacea (Troth et al., 2020); Chordata: Mammalia (Iso-Touru et al., 2021); Chordata: Reptilia (Farrell et al., 2022), whereas few have designed assays for targeting species belonging to multiple taxonomic groups (<5% of the studies) (Bronnenhuber and Wilson, 2013; Sepulveda et al., 2019; Thomsen et al., 2012; Wood et al., 2019b) (Fig. 4).

Most studies targeted a single species (approx. 69%) or 2 species (approx. 18%), while few have targeted more than 5 species (<5%) (Fig. 5A). A total of 404 different species have been targeted on these publications (**Table S2**) and the taxonomic groups for which more assays have been designed are Chordata: Actinopterygii (126 species), Chordata: Amphibia (78 species), Mollusca (44 species) and Arthropoda: Crustacea (34 species) (Fig. 5B).

Endangered species (which we considered species classified in the studies as Endangered, Critically Endangered, Vulnerable, Threatened, Imperilled, and of Special concern) have been the target of most studies in particular within Actinopterygii, Amphibia, Reptilia, Mammalia, and Mollusca (207 species) (Fig. 5B, **C**). The high focus on endangered species can be explained by the great interest in mapping and understanding their distribution, which is crucial for conservation management. In addition, most endangered species are often rare, with small population sizes and patchy distribution patterns; therefore, making these species suitable targets for the specific detection using eDNA-based assays. eDNA-based targeted detection can indeed be used as an important supplementary tool, especially in habitats and for species particularly challenging to survey (e.g., deep and turbid waters, large rivers and fast-flowing waters, aquatic species that burrow into the substrate, such as freshwater mussels or very elusive terrestrial animal species, such as wild cats or other carnivores) (Franklin et al., 2019; Parsons et al., 2018; Stoeckle et al., 2021; Strickland and Roberts, 2019; Sugiura et al., 2021; Williams et al., 2017) or where monitoring using classical methods are forbidden due to the possibility in leading to habitat alterations and/or destruction (Boon et al., 2019).

**Figure 5.**
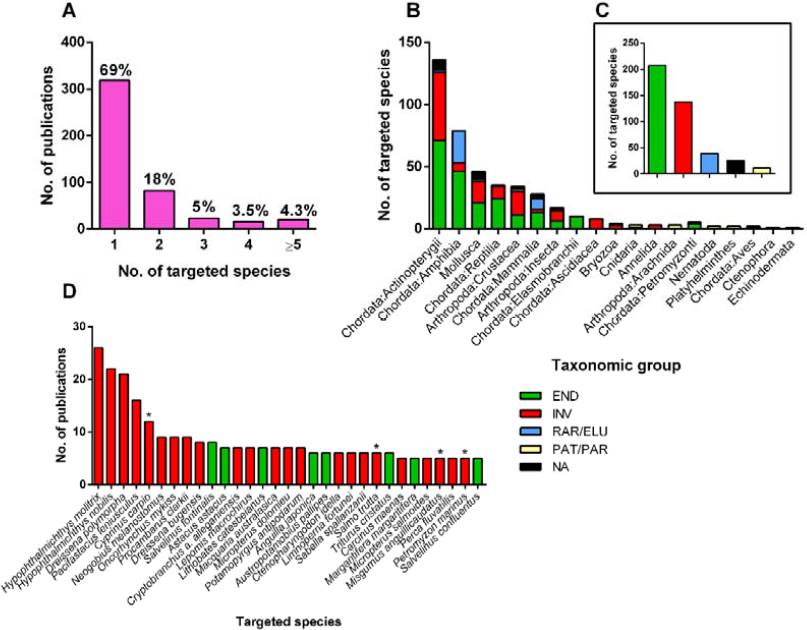
Number and % of publications *versus* the number of targeted species on each publication (A), number of targeted species and status per taxonomic group surveyed (B) and in general, in the 460 publications (C), and most targeted species (at least in 5 publications, approx. 1%) (D). * Species that have different statuses depending on the geographic region surveyed (e.g., endangered or invasive). END, endangered; INV, invasive; PAT/PAR, pathogen or parasite; RAR/ELU, rare or elusive; NA, not specified.

A high proportion of the species surveyed had the status of Invasive (137 species), in particular among the invertebrate taxonomic groups (Fig. 5B, **C**). Top-rank species that have been the focus of more studies using targeted assays are all invasive species in several regions of the world, in particular, the silver carp *Hypophthalmichthys molitrix* (Valenciennes, 1844) and the bighead carp *H. nobilis* J. Richardson, 1845, the zebra mussel *Dreissena polymorpha* Pallas, 1771, the signal crayfish *Pacifastacus leniusculus* Dana, 1852 and the Asian carp *Cyprinus carpio* Lineu, 1758, which have been the focus of more than 10, out of the 460 publications (Fig. 5D). *Cyprinus carpio* and *D. polymorpha* belong to the list of the 100 worst invasive alien species in the world (e.g., http://www.iucngisd.org/gisd/100_worst.php, accessed on 09^th^ February 2023), and other species, such as the silver and the bighead carps, are well known for provoking several negative ecological and economic impacts (e.g., in the Mississippi and Laurentian Great Lakes in North America).

On the other hand, among invertebrate taxa, a lower number of studies have been targeting pathogenic or parasitic animals, such as small crustaceans, cnidarians, nematodes, or Platyhelminthes (Fig. 5B, **C**). Micro-eukaryotic parasites are particularly challenging to detect and characterize due to their small size (typically <1 μm), as well as their intracellular or intra-organellar nature, and occurrence at low densities (Bass et al., 2015). DNA-based tools can indeed circumvent some of these barriers. However, they can also face challenges such as the fact of parasitic or pathogenic DNA being present in very small amounts in ecosystems, and in some cases, access to this DNA may require disruption of robust cysts or egg cases (Bass et al., 2015), probably explaining their lower adoption.

### 3.3. COI has been the most used genetic marker in the targeted detection of rare animal species

Most DNA markers in these targeted assays were designed to target regions within the mitochondrial genome (Fig. 6), namely the cytochrome *c* oxidase subunit I gene (COI) and the Cytochrome *b* gene (Cytb) in vertebrate animals (>300 cases) (Fig. 6A), while within invertebrate species there was a clear dominance of the use of the COI region for designing the assays (186 cases) (Fig. 6B). Group specificity of mitochondrial sequences, uniparental nature of inheritance, lack of recombination, relatively small genome, and a large number of copies in cells, make the mitochondrial genes well suited for analysing degraded genetic material (Ballard and Whitlock, 2004; Salas et al., 2007). In addition, oxidative processes that take place in the mitochondria and the lack of repair mechanisms lead to mutations in mitochondrial DNA. In particular, the sequences of the genes that code for elements of the respiratory chain, such as Cytb and COI, accumulate specific mutations which made them to vary considerably between species (Blaxter, 2003; Kumar et al., 2019; Linacre and Lee, 2016), which justifies their greater performance in targeted assays, in comparison to other markers.

Other commonly analysed mitochondrial DNA fragments include the ribosomal 12S RNA and 16S RNA genes and the control region or D-loop, which contain hypervariable regions (Habza-Kowalska et al., 2020) (Fig. 6A). In addition, we also found NADH genes to be widely used in particular for designing targeted assays for vertebrate animal species (Fig. 6A). These genes encode the NADH dehydrogenase subunits of respiratory complex I, that catalyse the oxidation of NADH by ubiquinone, and at least NADH2 has been useful to reveal genetic variation and diversity within bird species (Astuti and Prijono, 2016) (Fig. 6A).

**Figure 6.**
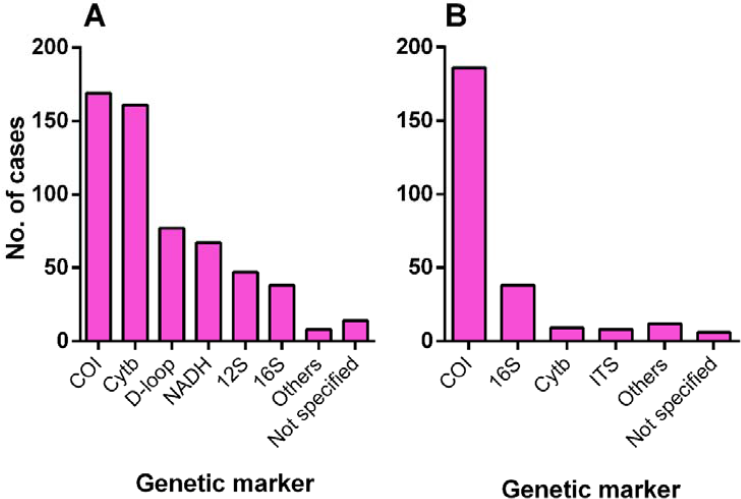
DNA markers that have been mostly employed in eDNA-based targeted detection of vertebrate (A) and invertebrate (B) animal rare species, in the 460 publications.

On the other hand, nuclear markers have been much less adopted (Fig. 6). In fact, eukaryotic cells can contain up to several hundred mitochondria, but only one nucleus, and thus, can carry thousands of copies of mitochondrial DNA *versus* only one nuclear genome with only two copies per nuclear gene. Among nuclear genes, the internal transcribed spacer region (ITS) of the nuclear ribosomal DNA has been employed for designing a few assays targeting invertebrate species (Fig. 6B). The ITS region contains two variable non-coding regions nested within the rDNA repeat, between the highly conserved small subunit, 5.8S, and the small and large subunits rRNA genes (Devi et al., 2022). In addition, the nuclear ribosomal DNA repeat is also available in thousands of copies in the nuclear genome, which facilitates its detection and amplification and is also a suitable target for the analysis of degraded genetic material. In particular, the ITS2 region has sufficient variability to distinguish closely related species (Yao et al., 2010). In the particular case of fish, nuclear DNA markers may be useful as an alternative or as an additional marker in species identification (Piggott, 2016) and potentially to delineate species boundaries and detect hybridization, when mitochondrial DNA markers are unable to do it (Hardy et al., 2011; Ward et al., 2009).

Most developed assays targeted small fragments (<200 bp, **Table S2**). Since eDNA is usually degraded, a short amplicon would increase the possibility of detection by PCR. In addition, in qPCR-based eDNA studies, which as indicated by our review is the most used platform (see below), the recommended amplicon size for a TaqMan probe is less than 150 bp. Amplicons that are too short may decrease PCR specificity, hence primer specificity needs to be well evaluated by testing against sequences from co-occurring and congeneric species (Meusnier et al., 2008). Increasing marker length may rise primer specificity, but to the detriment of amplification success (Harper et al., 2020; Valsecchi et al., 2022; Wei et al., 2018). For instance, Wei and co-authors (Wei et al., 2018) found that the qPCR copy number using a shorter marker was 12.1 times higher than the obtained using a longer marker within COI (126 bp *versus* 358 bp), targeting a benthic amphipod on sediment eDNA. Similar conclusions were reached by Harper and co-authors (Harper et al., 2020) when targeting green turtle eDNA in water (253 bp *versus* 488 bp, for D-loop) or by Valsecchi and co-authors (Valsecchi et al., 2022) (71 and 146 bp *versus* 216 bp, within 12S and 16S rRNA genes, respectively), for Mediterranean monk seal eDNA obtained from several sources. The use of small DNA fragments (90-120 bp) has been previously recommended to reach higher copy numbers (Rees et al., 2014; Saito and Doi, 2021), since eDNA degradation is accelerated in longer segments. On the other hand, no difference in performance was found when comparing 12 species-specific primer pairs producing amplicons from the Cytb gene of the Yangtze finless porpoise, ranging from 76 to 249 bp (Ma et al., 2016) or by Piggot (Piggott, 2016) (78-390 bp) on the detection of the endangered Macquarie perch in water eDNA. In the latter, since a closed system was surveyed (i.e., dams), the effect of the marker length may be smoother, contrarily to environments where eDNA can be more exposed to degradation, such as rivers or streams. In addition, for systems where organismal densities or biomasses are higher, the effect of marker length might also be lower (Piggott, 2016).

### 3.4. Quantitative PCR has been the most adopted platform in rare animal species detection

Three main platforms have been adopted in studies aiming at the targeted detection of rare animal species: quantitative PCR (qPCR), conventional PCR (cPCR), and digital droplet PCR (ddPCR). Quantitative PCR was used in more than half of the publications (>300 publications, approx. 73%), followed by conventional PCR, which has been employed in particular in earlier studies (Dejean et al., 2011; Ficetola et al., 2008; Goldberg et al., 2011; Jerde et al., 2011) and digital droplet PCR, adopted in more recent publications (Brand et al., 2022; Thalinger et al., 2019; Wood et al., 2020, 2019b) (Fig. 7). However, it should be noted that the period of availability of the ddPCR technology is much shorter than the other two platforms (only since 2011). Whereas cPCR based-assays strictly test for the presence or absence of a species, through the visualization of the expected PCR-amplicon band in an agarose gel, both qPCR and ddPCR provide also a quantitative estimate, enabling the quantification of the amount of the target species DNA in the environmental sample. In qPCR, the amplification products are continuously detected in the course of the reaction, due to the intercalation of a fluorescent dye or a specific probe labelled fluorescently (i.e., species-specific) in the amplification process. The amount of DNA is estimated through the use of a standard curve (using known amounts of target DNA), where the qPCR signal measurement is based on a Ct value (threshold cycle), corresponding to the point where the fluorescent signal exceeds a threshold. On the other hand, the microfluidics-based ddPCR consists of the partition of the PCR solution containing the DNA template into thousands of discrete droplets, where a PCR reaction occurs (Whale et al., 2012). Each droplet can contain either DNA molecules of the target (“1”) or not (“0”), which will lead respectively to the presence or absence of a fluorescent signal. After multiple cycles, samples are checked for fluorescence and the positive fraction recorded (the sum of all “1”) accurately indicates the initial amount of template DNA.

**Figure 7.**
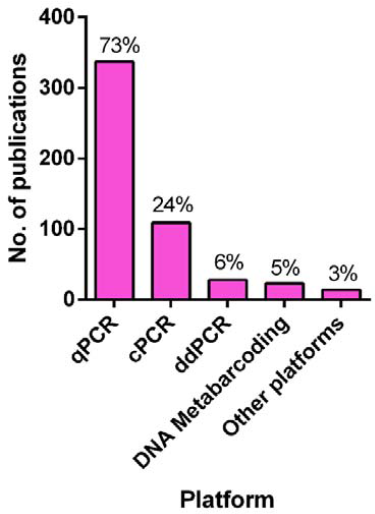
Platforms employed in eDNA-based targeted detection of rare animal species in the 460 publications examined.

In our review, we found that few studies have directly compared the sensitivity of the three platforms in the detection of rare animal species (Mauvisseau et al., 2019; Nathan et al., 2014; Qu and Stewart, 2019). While conventional PCR is the less sensitive platform, it has the advantage of being much cheaper, faster, and simpler, it can be carried out in any laboratory supplied with basic molecular biology equipment (i.e., a standard thermocycler, and a gel casting system) (Amberg et al., 2015; De Ventura et al., 2017; Nathan et al., 2014; Qu and Stewart, 2019; Valsecchi et al., 2022; Wilcox et al., 2013; Williams et al., 2017; Xia et al., 2018a). In addition, it may be sufficient, and reliable as well, in situations where researchers and/or environmental managers only require data on the presence or absence of target species or in places with limited infrastructures (De Ventura et al., 2017; Nathan et al., 2014). For instance, De Ventura and co-authors (De Ventura et al., 2017) found cPCR less prone to false positives and negatives (since it has lower sensitivity) than qPCR.

While both qPCR and ddPCR have been found to produce either similar estimates of DNA concentrations (Nathan et al., 2014) or strong relationships between marker copy numbers and abundances (Wood et al., 2019b), most studies found ddPCR more sensitive than qPCR. This may be because in ddPCR each sample is partitioned into 15,000–20,000 microfluidic droplets (see above), where the amplification reaction occurs independently and concentrations of PCR inhibitors can be strongly reduced (Banks et al., 2021; Brys et al., 2021; Doi et al., 2015; Hunter et al., 2017; Jerde et al., 2016; Mauvisseau et al., 2019; Williams et al., 2017; Wood et al., 2019a).

On the other hand, eDNA metabarcoding has been rarely employed in the targeted detection of rare animal species (Aylward et al., 2018; Balasingham et al., 2018; Crane et al., 2021; Marshall et al., 2022; Peterson et al., 2022; Rojahn et al., 2021; Stepien et al., 2019). In this approach, sequences belonging to multiple species are obtained from complex environmental samples via HTS, using short regions of one or a few marker genes, which are targeted with broad-spectrum or group-specific primers (Deiner et al., 2017; Taberlet et al., 2012a). Comparisons between metabarcoding and targeted approaches indicated that qPCR or cPCR or ddPCR are more sensitive in the detection of single species or a small set of species, in particular when the target DNA occurs at low densities (Banks et al., 2021; Blackman et al., 2020, 2018; Harper et al., 2018a; Moss et al., 2022; Roy et al., 2018). For instance, qPCR was highly effective in detecting the invasive tunicate *Didemnum vexillum* Kott, 2002 from seawater eDNA, whereas metabarcoding was unable to recover it, even at locations where it is known to be present, but detected several other established invasive species (Gargan et al., 2022). DNA metabarcoding prime application is for characterizing the taxonomic composition of whole communities, which can be particularly advantageous when several target species need to be simultaneously detected and identified (Roy et al., 2018; Thomsen et al., 2012), but it is less effective for sensitive and cost-effective screening of specific species (Roy et al., 2018). The choice between active *versus* passive surveillance for rare species depends on the study-specific aims. Active surveillance is highly sensitive in detecting rare DNA, while passive surveillance has the potential to identify unforeseen species, including early detection of invasive species. Therefore, employing a combination of active and passive surveillance using the same eDNA sample can provide significant advantages in invasive species management (Blackman et al., 2020; Simmons et al., 2016). Nevertheless, a few targeted metabarcoding assays have been already employed to simultaneously identify and distinguish closely related species [e.g., *D. polymorpha* and *D. rostriformis* (Deshayes, 1838)], as well as their phylogenetically-close relatives, and even to probe their population genetic structure across temporal and spatial scales (Marshall and Stepien, 2019).

Multiplex approaches have been less adopted but can be more cost-effective than DNA metabarcoding when the target is a few *a priori* well-known species (Jo et al., 2020; King et al., 2022; Robinson et al., 2018b, 2018a; Rodgers et al., 2020; Tsuji et al., 2018; Wozney and Wilson, 2017) (**Table S2**). The use of multiplex designs can allow the simultaneous detection of numerous species, reducing processing and handling times, as well as the risk of contamination, lowering costs and reducing the amount of DNA extract required for testing (Rodgers et al., 2020; Wozney and Wilson, 2017). For instance, Robinson and co-authors (Robinson et al., 2018a) developed a multiplex assay for the simultaneous detection of the invasive signal crayfish, the endangered white-clawed crayfish, and the crayfish plague pathogen using eDNA, allowing to assess of potential contributing factors to native crayfish decline with greater sensitivity, specificity, and efficiency than trapping, or single-species assays. Multiplex assays may also involve the use of different genetic markers, which may significantly improve the specificity of the assay and organism detection (Evans et al., 2016) or to better differentiate highly genetically similar species (Catanese et al., 2022). For example, Farrington and co-authors (Farrington et al., 2015) showed that the use of multiple highly sensitive markers maximized detection rates of the invasive silver and bighead carps, greatly improving the resolution of already implemented assays in eDNA-based surveillance programs.

### 3.5. Challenges for detecting rare animal species through eDNA-based targeted surveys

#### 3.5.1. Environmental factors effects

Most of the experiments dedicated to the targeted detection of rare animal species have been conducted in the field (>70%) (Fukumoto et al., 2015; Miralles et al., 2016; Sepulveda et al., 2019; Wood et al., 2019a) (Fig. 8, **Table S2**). On the other hand, only 17% of the studies were performed both in the field and also under controlled environments (i.e. mesocosms, aquariums, lab tanks, and artificial ponds, among others) (Dejean et al., 2011; Ito and Shibaike, 2021; Ladell et al., 2019; Matejusova et al., 2021; Mauvisseau et al., 2018; Mizumoto et al., 2018; Takeuchi et al., 2019; Troth et al., 2020; Turner et al., 2015; Yoshitake et al., 2019), and an even lower percentage was conducted exclusively under controlled conditions (8.7%) (Jerde et al., 2016; Mizumoto et al., 2018; Seymour et al., 2018; Stoeckle et al., 2017) (Fig. 8, **Table S2**). Experiments conducted under controlled conditions are extremely important to analyse in more detail: i) the amount and integrity of shed DNA; ii) dynamics of eDNA persistence, degradation, and transport, and iii) how taxa, sample type (i.e., water, soil or sediment) and ecosystem-specific factors (e.g., temperature, UV radiation, pH, presence of PCR inhibitors, salinity, among others) can affect i) and ii) (**Table S3**).

**Figure 8.**
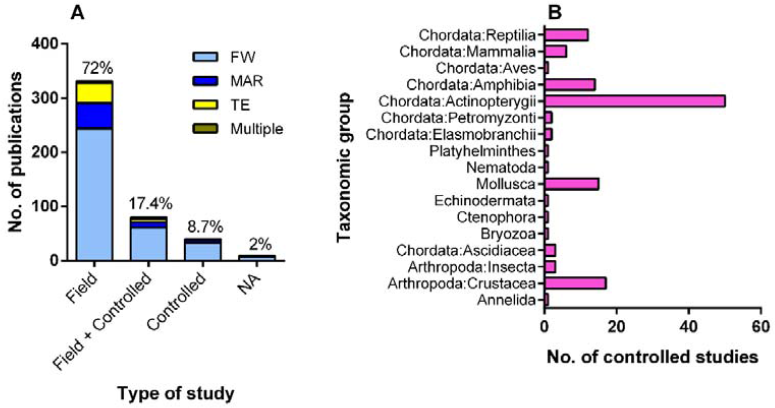
Type of study in publications using targeted assays for detecting rare animal species separated by the environment surveyed and % of publications using each type of study (numbers above bars) (A) and taxonomic groups that have been addressed in controlled studies (B). FW: freshwater ecosystems, MAR: marine ecosystems, TER: terrestrial ecosystems, MUL: multiple ecosystems, NA: not specified.

Few studies have directly measured how much eDNA an organism sheds into the environment over time (Klymus et al., 2015; Sassoubre et al., 2016; Thomsen et al., 2012), and most of these studies have been focussed on freshwater ecosystems (Fig. 8A) and fish species (Fig. 8B, **Table S3**). eDNA shedding rates have been found to depend on several factors (Fig. 9, **Table S3**), such as:

##### i) type of organism or characteristics of the species under analysis

(Goldberg et al., 2011; Thomsen et al., 2012), with animals with a hard or keratinized carapace or low-secretion taxa (e.g., reptiles, large invertebrates with exoskeletons or shells, mussels with closed valves) shedding less eDNA than animals with semipermeable skins or outer layers or higher-secretion taxa (e.g., amphibians, fish) (Adams et al., 2019; Danziger et al., 2022; Danziger and Frederich, 2022; Nordstrom et al., 2022), which might in part explain the higher adoption of these tools for the detection of fish and amphibians (Fig. 4 and 5). For instance, in controlled experiments, the water eDNA of the painted turtle was amplified only in the highest-density treatments, suggesting that detection in field samples using eDNA may be particularly difficult (Adams et al., 2019; Raemy and Ursenbacher, 2018).

##### ii) number/density or biomass of organisms

with numerous authors finding positive correlations between species biomass, density and detection probability and efficiency, e.g., fish (Brys et al., 2021; Dejean et al., 2011; Díaz-Ferguson, 2014; Mizumoto et al., 2018; Robinson et al., 2019; Sassoubre et al., 2016; Schloesser, 2018); amphibians (Dejean et al., 2011; Goldberg et al., 2011); reptiles (Adams et al., 2019; Tarof et al., 2021); molluscs (Blackman et al., 2020; Goldberg et al., 2013; Ito and Shibaike, 2021; Mauvisseau et al., 2017; Miralles et al., 2019; Xia et al., 2018b) or crustaceans (Baudry et al., 2021; Harper et al., 2018a); but with no correlations being also found in some other studies, e.g., reptiles (Raemy and Ursenbacher, 2018) and crustaceans (Danziger et al., 2022);

##### iii) organism size and developmental stage

with fish eDNA release rates being found to be higher in adults than in juveniles (Maruyama et al., 2014; Mizumoto et al., 2018), but no effect was found when eDNA concentration is adjusted taking into account total biomass (Mizumoto et al., 2018). For crustaceans, the presence of eggs increased eDNA concentrations per unit of mass (Crane et al., 2021; Dunn et al., 2017). In addition, the eDNA amount of a species can increase during its breeding period (Spear et al., 2015) and may vary considerably through time among individuals maintained under the same conditions (Klymus et al., 2015; Pilliod et al., 2014; Strickler et al., 2015);

##### iv) water temperature

in general, high temperatures have been reported to either produce higher eDNA shedding rates for fish species (35°C *versus* 23 and 29°C) (Robson et al., 2016) or to not have any effect (19 *versus* 25 *versus* 31°C) (Klymus et al., 2015);

##### v) other factors

that although less studied, have been shown to influence eDNA shedding rates. For instance, **exposure to stress** (Pilliod et al., 2014) or **feeding activity** (Klymus et al., 2015) have been found to increase eDNA shedding rates in amphibians and fish, respectively. Therefore, high eDNA shedding rates might be also found in seasons of higher nutrition (summer and spring). On the other hand, the presence of natural substances and substrates such as humic acids, sediments (Stoeckle et al., 2017), clay or topsoil (Buxton et al., 2017), the presence of other organisms (e.g., algae) (Stoeckle et al., 2017), filter-feeders (Mächler et al., 2018), the water pH (Tsuji et al., 2017) and current velocity (Malekian et al., 2018) can delay eDNA release or reduce the detection probability of the target organisms.

**Figure 9.**
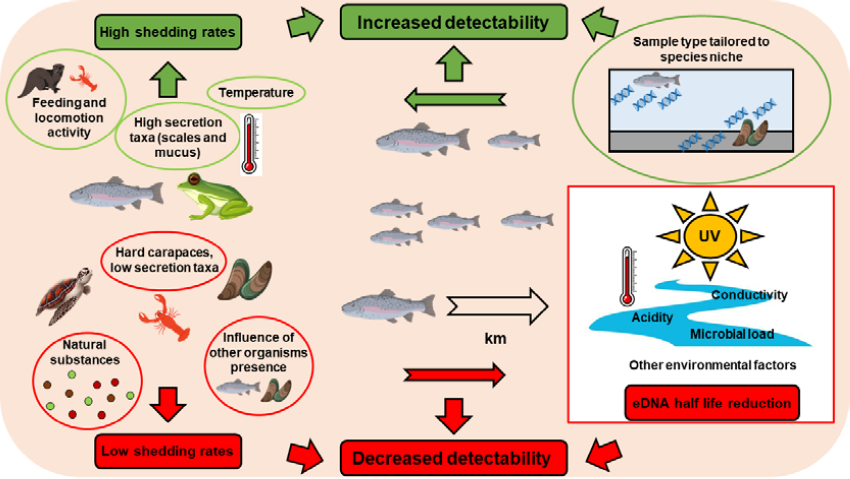
Factors affecting species detectability through eDNA.

Seymour and co-authors (Seymour et al., 2018) defined eDNA persistence dynamics as the relationship between physical, abiotic, or biotic factors and the degradation and localized detection of eDNA in natural ecosystems. In addition, detection probability depends on the ratio between the DNA released by the organism, and the DNA degraded by environmental factors (Dejean et al., 2011) and settled, such as temperature (Kasai et al., 2020; Lance et al., 2017; Nevers et al., 2018; Strickler et al., 2015), water conductivity and dissolved solids (Tarof et al., 2021); UV radiation (Day et al., 2019; Strickler et al., 2015), acidity (Seymour et al., 2018), microbial load (Lance et al., 2017), which have been found to reduce eDNA half-life and accelerate degradation (Fig. 9, **Table S3**). In addition, the environmental sample type chosen can also strongly influence species detectability. For instance, target eDNA concentrations have been found to be higher in sediments than in water (Kusanke et al., 2020; Nevers et al., 2020; Turner et al., 2015) (**Table S3**). In addition, sediment sampling may greatly increase the chances of detecting benthic-dwelling organisms, spending most of their life near the bottom of water bodies, e.g., Weatherfish (Kusanke et al., 2020) or sea lamprey larvae (BaltazarLSoares et al., 2022). However, eDNA has been found also to persist for longer periods in sediments, than DNA that is dissolved or suspended in water, mostly due to particle settling and/or retarded degradation of sediment-adsorbed DNA molecules (Turner et al., 2015). One way to increase the chances of benthic species detection may be to sample water near the bottom (Lor et al., 2020; Xia et al., 2018b), while surface sampling might be more adequate to detect species spending most of their life cycle at the surface (Moyer et al., 2014). In terrestrial samples, the choice of the sample type can be also very critical when dealing with rare species, probably due to a patchier distribution of eDNA. For instance, although eDNA from the terrestrial and small-bodied eastern red-backed salamander was positively detected in skin swabs and faecal samples, no eDNA was found in soil samples collected directly underneath wild-caught living salamanders (Walker et al., 2017).

Although in most controlled studies eDNA detectability has been found to decrease with time after removal of the source DNA species, the rate of subsequent eDNA decay might be highly variable among different animals: up to 7 days for the signal crayfish (Harper et al., 2018a); 21 to 44 days for the New Zealand mud snail (Goldberg et al., 2013); less than 1 month for the American bullfrog and the Siberian sturgeon (Dejean et al., 2011); at least 1 month for the Bighead carp (after carcasses deposition) (**Table S3**). However, eDNA persistence has been also shown to be highly dependent on organisms’ density (Dejean et al., 2011; Goldberg et al., 2013; Harper et al., 2018a). For instance, Harper and co-authors (Harper et al., 2018a) were able to detect signal crayfish DNA 7 days after organisms’ removal in high-density tanks, but only after 72 hours in low-density tanks. In addition, controlled studies indicated that positive eDNA detections can also be achieved with dead organisms: fish carcasses (Kamoroff and Goldberg, 2018); crustaceans’ carcasses (Curtis and Larson, 2020) or molluscs empty shells (Rasmussen et al., 2021), but the distance from the source seems to reduce the chances of detection, decreasing the probability of getting false positives (experiments with fish species in cages and with molluscs) (Dunker et al., 2016; Robinson et al., 2019; Xia et al., 2018a). The study by Blackman and co-authors (Blackman et al., 2020) found that the most significant predictor of quagga mussel DNA copy number and relative read count was the distance from the source population, even more than density. Even so, previous findings pointed out that for non-benthic species eDNA can be patchily distributed horizontally, even at a small spatial scale of tens to hundreds of meters (Eichmiller et al., 2014) and persist over relatively large distances from the established populations of the target organisms in natural river systems (up to 10 km for the cladoceran freshwater water flea) (Deiner and Altermatt, 2014). For example, Lamarie and co-authors (Laramie et al., 2015) did not find any consistent relationship between stream distance and eDNA concentrations of the chinook salmon. In addition, the transport of eDNA via predatory species (e.g., piscivorous birds) or deposition in slime residues and predator faeces can also be effective sources of eDNA, eventually leading to false positives in unpopulated habitats (Guilfoyle et al., 2017; Merkes et al., 2014).

In marine and terrestrial systems, controlled studies on eDNA persistence have been rarer. Marine systems present a set of features that differ from freshwaters, in what respects eDNA stability. Several studies indicate that eDNA degrades generally faster in marine systems (Sassoubre et al., 2016; Thomsen et al., 2012), however, DNA of the Mediterranean fanworm and the club tunicate was still detectable up to 94 hours, after organisms’ removal in controlled aquarium experiments (Wood et al., 2020). On the other hand, experiments with terrestrial snakes also demonstrated that eDNA declined up to approximately one week after organism removal (Kucherenko et al., 2018; Ratsch et al., 2020; Walker et al., 2017), but was still amplifiable after 7 days in controlled experiments with DNA from a terrestrial small bodied salamander (Walker et al., 2017).

#### 3.5.2. Closely related and co-occurring species, species hybridization and intra-genetic variation

The accuracy of the targeted assays for detecting rare animal species strongly depends on the availability of genetic data from closely related co-occurring species. Such data is crucial to assess assays’ genetic markers and primers’ specificity. Lack of sufficient specificity can result in both false positive and negative results, particularly in the presence of abundant and related species (Wilcox et al., 2014). In addition, closely-related species may hybridize (Antognazza et al., 2019; Farley et al., 2018; Fukumoto et al., 2015), which can further complicate the design of specific assays able to distinguish hybrids from non-hybrids, because of the maternal inheritance of mitochondrial DNA. This might be circumvented by using nuclear genes, but often the most popular nuclear targets currently available have an insufficient taxonomic resolution. The case is even more problematic when the target endemic species is closely related to an invasive exotic species (Fukumoto et al., 2015). For instance, a DNA-based survey for giant salamanders *Andrias japonicus* (Temminck, 1836) in the Katsura River basin performed by the Kyoto City Government in 2012, revealed that only 25 out of 125 captured individuals were pure endemic species, 6 were exotic [Chinese giant salamander, *Andrias davidianus* (Blanchard, 1871)], and 76 were hybrids (Fukumoto et al., 2015).

Designing highly species-specific primers for closely related species can be challenging since they can share high homology in the mitochondrial sequences. One way of increasing specificity is to use blocking primers (Wilcox et al., 2014). For instance, the addition of a blocking primer substantially increased assay specificity, without compromising sensitivity or quantification ability in a study using a purpose-designed TaqMan assay for eDNA detection of the endangered bull trout (*Salvelinus confluentus* Suckley, 1859) in the presence of the closely related and more abundant lake trout [*S. namaycush* (Walbaum, 1792)] (Wilcox et al., 2014) Other issues that can further complicate the design of species-specific eDNA assays are the intraspecific polymorphism and ambiguous taxonomic status of the target species (Dugal et al., 2022; Serrao et al., 2021; Utzeri et al., 2021; Wilcox et al., 2015; Yoshitake et al., 2019). Wilcox and co-authors (Wilcox et al., 2015) developed qPCR assays to distinguish westslope cutthroat trout [*Oncorhynchus clarkii lewsi* (G. Suckley, 1856)], Yellowstone cutthroat trout [*O. clarkii bouvieri* (Jordan & Gilbert, 1883)], and rainbow trout (*O. mykiss* Walbaum, 1792), which are of conservation interest both as native species and as invasive species across each other’s native ranges. The authors found that local polymorphisms within westslope cutthroat trout and rainbow trout posed a challenge to designing eDNA-based assays that are generally employed across the range of these widely-distributed species. In addition, in Europe, the existence of different genetic lineages within the Louisiana crawfish (Oficialdegui et al., 2019) was probably the main reason that led to the failure of eDNA probes to detect target populations in France (Mauvisseau et al., 2018; Tréguier et al., 2014) and that had previously worked well with the less variable Chinese populations (Cai et al., 2017). For instance, to take into account intra-specific genetic variability, Serrao and co-authors (Serrao et al., 2021) developed three assays to detect big brown bats eDNA for eastern, western, and southern North America and were highly successful in detecting very low concentrations of bat eDNA from air, water, and soil in different geographic regions. In addition, previously developed assays that work at a particular geographic location could be unsuitable for species detection at other places due to matches with the sequences of co-occurring species (Ogata et al., 2022). Thus, these case studies reveal that eDNA-based assays developed for a particular species may have to be refined taking into account site-specific genotypes.

## 4. Final considerations

As demonstrated by the current review, thanks to the high sensitivity in the detection of rare and elusive animal species, eDNA-based approaches evolved rapidly, and have been extensively applied for conservation and management purposes. Indeed, studies made so far have shown the great potential of eDNA-based species-specific detection to: i) increase and improve the data available on the presence/absence or occurrence of rare species (i.e., site occupancy), leading to a better understanding of present and historical patterns of species distribution (Boyd et al., 2020; Collins et al., 2019; Macgregor et al., 2021; Pitt et al., 2017; Sigsgaard et al., 2015; Tingley et al., 2019; Turner et al., 2015), ii) evaluate the success of restoration implementation efforts of endangered species (Budd et al., 2021; Feist et al., 2022; Goldberg et al., 2018; Hempel et al., 2020; Hossack et al., 2022; Kamoroff & Goldberg, 2018; Wineland et al., 2019), iii) early detect non-indigenous and invasive species (Créach et al., 2022; Koel et al., 2020; Sepulveda et al., 2019), iv) confirm eradication of invaders, namely where positive eDNA detections can trigger more in-depth sampling to find invasive specimens and remove them before native species reintroductions, or to postpone native species re-introductions, while invasives are still in place (Bylemans et al., 2016; Carim et al., 2020; Dunker et al., 2016; Furlan et al., 2019; García-Díaz et al., 2017; Miralles et al., 2016; Robinson et al., 2019; Schumer et al., 2019); v) construct exclusion barriers to prevent invasives spread (Bylemans et al., 2016; Carim et al., 2020; Hunter et al., 2019; Miralles et al., 2016) and vi) better estimate range limits, of both endangered or invasive species (e.g. Gargan et al., 2022; Rose et al., 2019; Westhoff et al., 2022). However, it remains less clear how results have been translated into management actions (Sepulveda et al., 2019), but for some species eDNA-based detection is already in place, aiding in decision-making (Biggs et al., 2015; Laramie et al., 2015). For instance, DNA-based protocols have been employed as a trigger in the surveillance of the bighead and silver carps *H. nobilis* and *H. molitrix*, in the Great Lakes region (USA and Canada), where positive eDNA detections that follow a standard and very rigorous operating procedure prompt intensive molecular and nonmolecular monitoring to locate the fish populations. In addition, the great crested newt *Triturus cristatus* (Laurenti, 1768) is the first species to be routinely monitored using eDNA (approved by Natural England in 2014), with the specific assay being offered as a commercial service by several ecological consultancies in the UK. Indeed, developers can even be prohibited from interventions in wetlands where there have been positive eDNA detections of the great crested newt (Harper et al., 2018b).

The relatively large proportion of methodological development studies we recorded, highlights the suboptimal status of many assays and reinforces the continuing need for further adjustment, validation, and optimization of eDNA techniques, from sampling, through laboratory protocols and up to data analyses. Among other precautions, it is fundamental to take into account the particular characteristics of the target species and survey sites since the dynamics of eDNA might differ drastically among taxa, study systems, and across climatic zones, and therefore, thinly customized system-specific assays are required (Harper et al., 2019; Sales et al., 2021). In addition, beyond testing specificity, sensitivity (minimum eDNA concentration required for the species to be detected) of newly developed assays must also be optimized and verified in natural conditions to reduce the detection uncertainty. However, in a recent review Xia and co-authors (Xia et al., 2021) found that for most studies using newly designed markers (82.4%), researchers do not screen their chosen markers for sensitivity, with almost half of the studies not reporting the limit of detection of the assays. Indeed, for legal implementation end-users need to recognize the power and limitations of existing tools, to know how eDNA-based targeted detection works, what are the limitations, and what can offer to environmental managers in comparison with wellLestablished monitoring methods (Darling, 2019). It is also vital that managers know how to use the tool in the most appropriate way, how to interpret the results, and how these can influence decisions. To this end, the availability of well-established manuals on best practices and decision-support frameworks, that account for error minimization and quantification (Bruce et al., 2021; Darling, 2019; Sepulveda et al., 2020), will contribute to higher adoption and implementation of these tools in regular monitoring, and ultimately for more accurate monitoring and conservation of rare animal species.

## Supporting information

All supplementary material (Tables + figures)

## Acknowledgements

This work was funded by the project “River2Ocean – Socio-ecological and biotechnological solutions for the conservation and valorization of aquatic biodiversity in the Minho Region” (NORTE-01-0145-FEDER-000068), co-financed by the European Regional Development Fund (ERDF), through Programa Operacional Regional do Norte (NORTE 2020) and by the “Contrato-Programa” UIDB/04050/2020, funded by national funds through the Foundation for Science and Technology (FCT I.P). Financial support granted by the FCT to SD (CEECIND/00667/2017) and by the project ATLANTIDA (NORTE-01-0145-FEDER-000040), funded by Programa Operacional Regional do Norte (NORTE2020) to LS, is also acknowledged.

